# Phaeochromocytomas and paragangliomas harbour tumour-initiating SOX2+ stem cells

**DOI:** 10.1101/2025.07.03.663006

**Authors:** Yasmine Kemkem, Mark Quinn, Bence Kövér, Alice Santambrogio, James Kaufman-Cook, Olivia Sherwin, Laura D. Scriba, Dimitria Brempou, Miriam Vazquez Segoviano, Dylan Cameron, Ilona Berger, Wen Ng, Daisuke Nonaka, Marily Theodoropoulou, Christina Pamporaki, Paul V. Carroll, Louise Izatt, J Paul Chapple, Stefan R. Bornstein, Nicole Bechmann, Charlotte Steenblock, Rebecca J. Oakey, Cynthia L. Andoniadou

**Affiliations:** Centre for Craniofacial and Regenerative Biology, King’s College London, London, SE1 9RT, United Kingdom; Department of Medical and Molecular Genetics, King’s College London, SE1 9RT, United Kingdom; Department of Internal Medicine III, University Hospital Carl Gustav Carus, Technische Universität Dresden, Fetscherstrasse 74, 01307 Dresden, Germany; Guy’s & St. Thomas’ NHS Foundation Trust, London, SE1 9RT, United Kingdom; School of Cancer & Pharmaceutical Sciences, King’s College London, London SE1 9RT, United Kingdom; Medizinische Klinik und Poliklinik IV, LMU Klinikum, LMU München, Munich 80336, Germany; Department of Clinical Genetics, Guy’s and St Thomas’ NHS Foundation Trust, London, SE1 9RT, United Kingdom; William Harvey Research Institute, Faculty of Medicine and Dentistry, Queen Mary University of London, London EC1M 6BQ, United Kingdom

**Keywords:** PPGL, PCC, PGL, Stem cells, SOX2, sustentacular cells, tumour-initiating cells, cancer stem cells, phaeochromocytoma, paraganglioma

## Abstract

Phaeochromocytomas (PCCs) and paragangliomas (PGLs), are rare neuroendocrine tumours that arise in the neural crest (NC)-derived adrenal medulla and the paraganglia, respectively. Approximately 10%-15% of patients with PCCs and 35%-40% with PGLs go on to develop metastatic disease, leading to a reported median overall survival of 7 years. The development of prognostic markers and subsequent personal therapeutic strategies are hindered by a lack of understanding of tumourigenesis. In other organs, cells with stem-like properties are at the root of tumour initiation and maintenance, due to their ability to self-renew and give rise to differentiated cells. We have recently shown that, in the human adrenal, a subset of sustentacular cells, endowed with a support role, are in fact SOX2+ postnatal adrenomedullary stem cells, that are specified along the neural crest migratory route. In this study, we intended to determine if SOX2+ cells in PCCs and PGLs can behave as tumour-initiating stem cells. Using expression and transcriptomic studies, we demonstrate the presence of SOX2/*SOX2*-expressing cells across a broad range of PCCs and PGLs, irrespective of tumour aggressiveness, location, and causative mutation. *In silico* analyses reveal the co-expression of SOX2 and chromaffin cell markers in the tumour, and the active proliferation of these double-positive cells. Isolation of these cells *in vitro* in stem cell-promoting media, and their xenotransplantation on chicken chorioallantoic membranes, demonstrates that they have the potential to expand and metastasise *in ovo*, supporting their potential as tumour-initiating cells.

## INTRODUCTION

Phaeochromocytomas (PCC) and paragangliomas (PGL), collectively known as PPGL, are adrenal and extra-adrenal neuroendocrine tumours, respectively, composed of adrenaline- and noradrenaline-secreting chromaffin cells. They arise from neural crest-derived cells of the sympathetic and parasympathetic nervous system and according to WHO 2022 5th edition, all PPGLs have metastatic potential (1). With scarce therapeutic approaches available, the median survival rate of patients with metastatic PPGLs is ∼7 years (2). Even PPGLs classified as benign are associated with high morbidity and mortality due to comorbidities related to excessive catecholamine production and secretion, that include hypertension, arrhythmia and stroke (3, 4).

PPGLs are associated with the highest degree of heritability in human neoplasms, with approximately 40% carrying germline mutations or deletions in at least one of several susceptibility genes that include *VHL*, *SDHA*, *SDHB*, *SDHC*, *SDHD*, *SDHAF2, RET*, *NF1*, *TMEM127*, *MAX*, *H3F3A* and *HIF2A* (5). The most frequent cause of inherited PPGL are germline pathogenic variants in genes encoding for subunits of the mitochondrial succinate dehydrogenase (SDH) enzyme complex: *SDHA, SDHB*, *SDHC* and *SDHD* (6). Patients with *SDHB*-mutant PPGLs present with higher morbidity and mortality (7–9). Pathogenic germline *SDHB* variants have been detected in 42% of metastatic PPGLs (reaching 80% in paediatric metastatic PPGLs), but in a multivariate analysis on adult patients, they did not act as a prognostic factor of overall survival (2). In *SDHB*-mutant PPGLs secondary driver events in *TERT* or *ATRX* have been associated with metastatic disease (10). Combined germline and somatic driver genetic events are known in over 70% of cases, enabling the classification of PPGL into three clusters: cluster 1 (pseudohypoxia), cluster 2 (kinase signalling) and cluster 3 (WNT signalling) (11).

Despite our extensive knowledge of PPGL genetics, the exact cell-of-origin of PPGLs is currently unknown, and this together with their heterogeneity, severely limits the generation of *in vivo* experimental models and the road to targeted therapies. The possibility of a cancer stem cell (CSC) or tumour-initiating cell (TIC) has been proposed but not demonstrated for these tumours (reviewed in (12)). A previous study reported the expression of SOX2, a transcription factor associated with stem cells of multiple tissues, in 12% of PPGLs analysed by immunostaining on tumour microarrays (13). The study did not provide data to support stem cell function in these tumours. We recently identified a postnatal population of SOX2+ stem cells in the normal mouse adrenal medulla, which derives from the migratory neural crest population during embryonic development and which colonises the paraganglia and the adrenal medulla (14). That work demonstrated that SOX2+ adrenomedullary stem cells contribute to the generation of new chromaffin cells throughout life in mice, and are specified from neural crest-derived progenitor cells as a distinct population. Confirmation of their presence in normal human adrenals, coupled with the location of PPGLs along sites populated by the neural crest, raises the possibility that neural crest-derived SOX2+ cells might act as stem cells in PPGLs. In this study, we set out to determine if SOX2+ cells of PPGLs can behave as cancer stem cells and candidate cells-of-origin of these tumours.

## MATERIALS AND METHODS

### Ethical Approval

All animal studies were performed under compliance with the Animals (Scientific Procedures) Act 1986, Home Office License P8D5E2773 (chicken), as well as KCL Biological Safety approval for project ‘Function and Regulation of Adrenal Stem Cells in Mammals’. Foetal adrenals were received from the Human Development Biological Resource (HDBR)/project ID 200587. PPGL tumour studies were conducted under King’s College London and were approved by the Cornwall and Plymouth Health Research Authority, study title: Investigation of genetic and epigenetic marks in cancer, IRAS project ID: 216133, REC reference: 17/SW/0018. Formalin-fixed, paraffin-embedded (FFPE) tumour and adrenal tissue samples with pathologically confirmed PPGLs were prepared by the Tumour and Normal Tissue Bank (TNTB) of the BioBank Dresden, originate from the archive of the Institute of Pathology (EK 59032007) of the University Hospital Dresden. Patients were included from the Registry and Repository of biological samples of the European Network for the Study of Adrenal Tumours (ENS@T, EK 407122010), the Prospective Monoamine-producing Tumour study (PMT study; EK 189062010) and/or PROSPHEO (NCT03344016; EK 210052017), all of which had ethical approval at the University Hospital Carl Gustav Carus Dresden. Informed consent was obtained from all patients. Additional FFPE tumour samples were obtained from Queen Mary under study title: Genetics of Endocrine Tumours (Cambridgeshire Research Ethics Committee Reference MREC 06/Q0104/133. Normal adrenal human tissue samples were obtained from the University Hospital Würzburg (Germany). Normal adrenal glands were collected as part of tumour nephrectomy and proven to be histologically normal. Informed consent was obtained for all samples; patients were not compensated for participation.

### Chorioallantoic membrane (CAM) assays

Fertilised Shaver Brown eggs were purchased from Medeggs Ltd.Developmental. Day 0 corresponds to the day eggs were placed in an egg incubator set at 37.8°C/60% humidity. Four days later, eggs were windowed using curved spring scissors, exposing the CAM. Windows were sealed and eggs placed in the incubator until day 10, when isolated stem cells were seeded onto the CAM. 25µl of a single-cell suspension (100,000 cells) were pipetted onto the CAM, within a 6mm diameter silicone ring. Eggs were sealed and placed back in the incubator until collection on Day 14. Chicks were killed by Schedule 1 method. Resected CAM and dissected chick’s lungs were washed in PBS and fixed accordingly.

### Primary culture of adrenomedullary and PPGL-derived stem cells

PPGL tumours and adrenal glands were dissociated using an enzymatic digestion solution of 50μg/ml DNAse I (Sigma, CAT#D5025), 10mg/ml Collagenase II (Worthington, CAT#LS004177), 2.5μg/ml Fungizone (Gibco, CAT#15290026), 0.1X Trypsin-EDTA (Sigma, CAT#59418C) in 1X Hank’s Balanced Salt Solution (HBSS) (Gibco, CAT#14025050). Samples were sequentially incubated at 37°C and manually triturated, until reaching a single-cell suspension. Dissociation enzymes were deactivated with serum-rich base medium (DMEM/F-12 (Gibco, CAT#31330-038) + 5% FBS (Merk, CAT#F0804) + 50μl/ml Penicillin-Streptomycin (Gibco, CAT#15070063). The cell suspension was centrifuged, and the pellet was resuspended and plated in stem-cell promoting media: base media + 20ng/ml bFGF (R&D Systems, CAT#234-FSE) + 50μg/ml cholera toxin (Sigma, CAT#C8052). For immunostaining, cells were plated on glass coverslips coated with 0.1% gelatine diluted in PBS. For CAM assays, stem cell colonies were expanded in stem-cell promoting media. On transplantation day, cells were trypsinised, labelled with a green-fluorescent chloromethyl derivative of fluorescein diacetate (CMFDA) intracellularly-activated CellTracker dye (ThermoFisher, CAT#C7025) as per the manufacturer’s instructions. Labelled cells were resuspended in stem-cell promoting media as a single-cell suspension for seeding onto CAM.

### Tissue processing for immunofluorescence and immunohistochemistry

For paraffin-embedding, resected grafts/CAMs were fixed in 10% neutral buffered formalin (NBF) (Sigma, CAT#HT501128) overnight at room temperature, on a roller. The following day, samples were washed and gradually dehydrated using ethanol series. Paraffin-embedded samples were sectioned at 5μm thickness. Chicks’ lungs were cryo-embedded. For this, they were fixed in 4% PFA at 4°C overnight. Samples were washed and placed in a cryoprotective 30% Sucrose/PBS solution overnight at 4°C. Samples were embedded in Optical Cutting Temperature compound (VWR, 361603E), flash-frozen, and cryo-sectioned at 15μm thickness.

### Immunofluorescence and immunohistochemistry

Paraffin sections were de-paraffinised with Neoclear and were gradually rehydrated in decreasing concentrations of ethanol. Antigen retrieval was performed in a decloaking chamber NXGEN (Menarini Diagnostics, CAT#DC2012-220V) at 110°C for 3 min using Dako Target Retrieval Solution, pH 9.0 (Agilent, CAT#S236784-2), as per the supplier’s instructions.

Immunohistochemistry was performed using the ImmPRESS Excel Amplified HRP Polymer Staining Kit Anti-Rabbit IgG (Vector Laboratories, CAT#MP-7602-50) as per the supplier’s instructions, using a rabbit anti-SOX2 antibody (Abcam, CAT#ab92494; 1:500). Nuclei were counterstained with Vector Haematoxylin QS (Vector Laboratories, CAT#H-3404-100) and slides were mounted in VectaMount Permanent Mounting Medium (Vector Laboratories, CAT#H-5000-60).

Immunofluorescence staining was performed as follows: sections were blocked at room temperature for 1h in a 10% Donkey Serum Blocking Buffer (0.15% glycine, 2 mg/ml BSA, 0.1% Triton X-100 in PBS). Sections were then incubated overnight at 4°C with primary antibody diluted in 1% Donkey serum/Blocking Buffer (mouse anti-HNA (Merck, CAT#MAB4383, 1:300). Following three five-minute washes in PBS + 0.1% Triton X-100 (PBST), sections were incubated in secondary fluorophore-conjugated antibodies (Goat anti-mouse594 (Abcam, CAT#ab150116, 1:500) and DAPI (Abcam, CAT# ab228549, 1:5,000) in 1% Donkey Serum/Blocking Buffer.

Cryosections were air-dried and washed in PBS. Fixed cells were washed in PBS. In both cases, samples were blocked at room temperature for 1h in Blocking Buffer (1% BSA, 0.1% Triton X-100, 5% goat serum), followed by a 1-hour incubation in primary antibodies diluted in 1% Goat Serum/Blocking Buffer (mouse anti-Tyrosine Hydroxylase (BD Biosciences, CAT# 612300, 1:500 – goat anti-SOX2 (RD Systems, CAT# AF2018, 1:500)), and a 1-hour incubation in fluorophore-conjugated secondary antibodies (Biotinylated Goat anti-mouse (Abcam, CAT# Abcam ab6788, 1:500), Donkey anti-goat647 (Abcam, CAT# Abcam ab150131, 1:500). For tyrosine hydroxylase staining, a biotin-streptavidin amplification was performed. Following secondary biotinylated antibody incubation, sections were washed in PBST and incubated at room temperature for 1h with fluorescent-labelled streptavidin (Life Technologies, CAT#S32355, 1:500). DAPI (Abcam, CAT# ab228549, 1:5,000) was added in the last incubation. Samples were washed and mounted in Vectashield Antifade Mounting Medium (Vector Laboratories, CAT#H- 1000-10).

### Imaging

Sections with immunohistochemistry staining were scanned using a Nanozoomer-XR Digital slide scanner (Hamamatsu). High magnification images were acquired with an Olympus BX34F Brightfield microscope. Cell culture images were obtained with an Olympus Phase Contrast microscope. Immunofluorescence imaging was performed using a Zeiss LSM980 confocal microscope (Zeiss Plan- Apochromat 20×/0.8 dry objective). Z-stacks were acquired with a 0.75μm step. Imaging files were processed with Fiji and Nanozoomer Digital Pathology view. Figures were created in Adobe Photoshop version 26.6.0.

### Computational studies

#### scRNA-seq and snRNA-seq dataset acquisition

PPGL tumour samples were obtained directly from the Guy’s and St Thomas’ NHS Foundation Trust (GSTFT) operating theatres. Patients were enrolled to this study through the departments of endocrinology and clinical genetics in GSTFT. All patients enrolled provided written, informed consent which granted access to tumour tissues and clinical data. Following resection, tumour samples were immediately reviewed and dissected by an expert clinical pathologist. Confirmed PPGL tumour samples were then transferred to our laboratory where they were either snap frozen and stored at -80°C or they underwent dissociation in a single-cell suspension in expectation of immediate scRNA-sequencing. Single- cell dissociation was performed using an enzymatic digestion of 10µl/ml DNAse I (Sigma, CAT#D5025), 20µl/ml FBS (Thermo Fisher Scientific, CAT #10270106), 10mg/ml Collagenase II (Worthington, CAT#LS004177), 1mg/ml Dispase-II (Merck, CAT#SCM133) and 40U/ml RNaseOUT (Thermo Fisher Scientific, CAT#10777019) in 1ml of 1X Hank’s Balanced Salt Solution (HBSS) (Gibco, CAT#14025050). Samples were sequentially incubated at 37 °C and intermittently mechanically dissociated and vortexed throughout. Samples were then passed through a 40µm filter (Cole-Parmer, CAT# UY-06336-63) and centrifuged. The enzyme dissociation mix was removed, and the pellet was washed with and eventually resuspended in HBSS and 5% FBS (Thermo Fisher Scientific, CAT #10270106). The single-cell suspension was then inspected under the microscope using Trypan Blue Solution, 0.4% (Thermo Fisher Scientific, CAT#15250061) for cellular number and viability. Frozen samples underwent nuclei extraction as per 10X protocols. Samples were kept at 4°C throughout. 3-50mg of frozen PPGL tumour tissue was transferred to a pre-chilled Sample Dissociation Tube (10X, CAT#2000564) with 200μl of Lysis Buffer (Lysis Reagent (10X, CAT#200558), Reducing Agent B (10X, CAT#200087) and Surfactant A (10X, CAT#2000559). Samples were dissociated mechanically and 300μl of Lysis Buffer was added. The samples were incubated on ice for 8 minutes. Dissociated tissues were then centrifuged through a pre-chilled Nuclei Isolation Column (10X, CAT#2000562). Flowthrough was vortexed and centrifuged, and the pellet was resuspended with 500μl of Debris Removal Buffer (10X, CAT#200560). Samples were centrifuged and supernatant was removed before the pellet was resuspended in 1ml of Wash and Resuspension Buffer (1X PBS, 10% BSA and RNase Inhibitor (10X, CAT#2000565)). This was repeated until once depending on debris, with the final pellet resuspended in 50-500μl Wash and Resuspension Buffer. The suspension was then inspected for nuclei number, viability, debris and clumping using Trypan Blue Solution, 0.4%. Single cells/nuclei were processed on the Chromium iX instrument using 3’ v3.1 gene expression profiling reagents (10x Genomics) to generate GEMs (gel bead-in-emulsion) for cell barcoding. Cells were loaded on the chip with an aim to capture 8,000-10,000 cells per sample. Full length cDNA was generated from poly-adenylated mRNA and amplified. Dual indexed sequencing libraries were prepared from the amplified cDNA and final libraries were evaluated on the Agilent TapeStation 4200 using a High Sensitivity D5000 ScreenTape (Agilent Technologies). The libraries were then pooled and sequenced on NextSeq 2000 (Illumina) at a depth of approximately 20,000 read pairs per cell.

#### Pre-processing for samples collected in this study

Samples were aligned to the GRCh38 reference genome using CellRanger 9.0.1 (10X Genomics) with default settings, meaning that intronic reads were also included for both single-cell and single-nucleus samples.

#### Additional publicly available samples

*SDHB* PPGL single-nucleus RNA-seq samples were accessed from the corresponding Figshare of Flynn et al. (doi.org/10.1038/s41467-025-57595-y) https://doi.org/10.6084/m9.figshare.25792479. The Figshare folder contained the CellRanger output matrices, from which the raw_feature_barcode matrix was used as input. Patient metadata for the corresponding samples was also accessed from the supplementary material of Flynn et al. (doi.org/10.1038/s41467-025-57595-y). Additional datasets for healthy adrenal medulla samples were accessed from the Figshare of Flynn et al. (doi.org/10.1038/s41467-025-57595-y) as well, though these are originally from Zethoven et al. (doi.org/10.1038/s41467-022-34011-3). Dataset identifiers for samples accessed from other publications were kept as original throughout.

#### Quality control (QC) and further processing

To separate true cells from cell-free droplets, we first filtered out barcodes with less than 800 UMIs. Percentages of ribosomal, and mitochondrial counts were determined using the scanpy (10.1186/s13059-017-1382-0) calculate_qc_metrics function with “RPS”, “RPL”, or “MT-” flags.

QC was then performed within scanpy using median +/- X * median absolute deviation filters, with x ranging from 4-5 for different metrics. Specifically, these included: percentage of mitochondrial counts (max = 25%, X=5), percentage of ribosomal counts (max = 30%, X=5), percentage of counts in top 20 genes (X=5), log1p total counts (X=4), log1p genes detected (X=4). Following this, doublets were identified and removed using the scrublet algorithm implemented in scanpy with default settings (10.1186/s13059-017- 1382-0).

#### Dataset integration

To jointly analyse single-cells/-nuclei from all transcriptomic datasets, we used the scVI variational auto- encoder framework (doi.org/10.1038/s41592-018-0229-2). Specifically, we set each patient ID (one per dataset) as the batch_key and set the percentage of mitochondrial (pct_mito) and ribosomal (pct_ribo) counts as continuous covariates. The model was set up using 2 hidden layers and a latent space with 30 dimensions. The generative end of the model was specified to be negative binomial. The 30-dimensional latent space was then used for nearest-neighbor identification, Leiden clustering (resolution=0.4, flavor=”igraph”) and UMAP visualisations, all with scanpy implementations (10.1186/s13059-017-1382- 0).

#### Cell type annotation

Leiden clusters of cells were annotated using established markers from previous work on PPGLs (doi.org/10.1038/s41467-025-57595-y; doi.org/10.1038/s41467-022-34011-3) as follows: Chromaffin_cells: HAND2, SLC18A1, TH, PHOX2A; Cortex_cells: CYP11B1, STAR; Sustentacular_cells: SOX10, CDH19, S100B, SOX2; T_cells: CD4, TRBC1, TRBC2; Macrophages: CSF1R, CD163, C1QA; Mast_cells: TPSAB1, TPSB2; B_cells: JCHAIN, CD79A, CD79B; Endothelial_cells: ROBO4, FLT1; Fibroblasts: COL1A1, COL1A2, PDGFRB. Following initial overclustering, clusters with the same broad identities were merged.

#### Pseudobulk differential expression analysis

To perform differential expression analysis, pseudobulk samples were generated from each cell type in each sample (where the cell type exceeded 30 cells) using the decoupleR package (doi.org/10.1093/bioadv/vbac016). In the DE analysis, only cell types with at least 3 pseudobulk samples were kept. We then used the Limma-voom workflow (DOI: 10.1186/gb-2014-15-2-r29) to find cell type specific markers, with the model: “∼ cell_type”, and contrasts between all cell-type pairs. Markers were assigned if they were significant in all comparisons. We applied a similar workflow to find differentitally expressed genes in metastasis and *SOX2* status, with the models “∼ 0 + cell_type + cell_type:Metastasis" and “∼ 0 + cell_type + cell_type:Chrom_SOX2_status".

#### Ligand-receptor interaction predictions

Ligand-receptor interaction prediction was performed using LIANA+ (https://doi.org/10.1038/s41556-024-01469-w). This approach was chosen as it allows the use of an ensemble of ligand-receptor algorithms, as well as an integrated ligand-receptor interaction database. Here, 5 algorithms (CellPhoneDB with 1000 permutations, Connectome, log2FC, NATMI, SingleCellSignalR) were used as implemented in the aggregate_rank function using a merged version of databases (called “consensus” in LIANA+).

#### LIANA+ signalling dotplots

Top ligand-receptor interactions were visualised using the native plotting functions within the LIANA+ package, using the commands li.pl.dotplot with the following arguments: colour=’magnitude_rank’, size=’specificity_rank’, inverse_size=True, inverse_colour=True, top_n=20, orderby_ascending=True. We have plotted interactions with Chromaffin cells as source, ranking by specificity. These were controlled through the “orderby”, “target_labels” and “source_labels” arguments.

#### Heatmap plotting

Heatmaps were generated following pseudobulking of datasets with decoupleR and counts per million (CPM) and log1p normalising the resulting anndata objects. Heatmaps were then generated with the scanpy sc.pl.heatmap function with row-wise (e.g. per gene) min-max normalised values, with the arguments log=False, standard_scale="var".

#### UMAP plotting

UMAPs were plotted using the scanpy function sc.pl.umap with the sort_order=True argument for featureplots to prevent the few expressing cells from being buried under the large number of non- expressing cells.

#### Annotation of top TFs and signaling genes

A list of all human TFs was accessed from "https://humantfs.ccbr.utoronto.ca/download/v_1.01/TF_names_v_1.01.txt” and the LIANA “consensus” database was used to retrieve gene names relevant to cell-cell signaling (either ligands or receptors in the database).

#### Code and data accessibility

Our code and processed data are available on https://github.com/Andoniadou-Lab/SOX2_PPGL. We provide a Jupyter Notebook of all Python code used for processing the datasets from CellRanger outputs. In addition, we provide an R script that was used for differential gene expression analysis with Limma- voom. Finally, we provide the cell type annotated AnnData object as a resource, allowing others to reproduce our work. Cell Ranger output files (raw and filtered count matrices) have been deposited on Figshare with the following DOI: 10.6084/m9.figshare.29447750.

## RESULTS

### SOX2+ cells are found in both phaeochromocytomas and paragangliomas, irrespective of mutation and tumour site

To determine the presence of putative stem cells in PPGL tumours, we focused on SOX2, which we previously established as a marker of progenitor/stem cells of the adrenomedullary lineage (14). Extending the previous report of stem cell markers in PPGLs (13), we tested for SOX2 expression on FFPE sections from a cohort of 19 PPGLs across different tumour sites, mutations and malignancy status. This cohort was composed of 12 adrenal PCCs and seven extra-adrenal PGLs, of which two were located in the carotid body, three abdominal, and two thoracic. Germline mutations in known genes were present in 15 patients, spanning *NF1*, *SDHC*, *RET* and *SDHB*, whilst three patients had somatic mutations in *RET* and *VHL*. Five of the tumours were metastatic, three of which harboured mutations in *SDHB*. Tumour data are found in **Table 1**. Immunostaining using antibodies against SOX2 identified positive cells in all tumours tested (**Figure 1 A-G**). Heterogeneity was seen across different tumour locations. SOX2-positive cells ranged from 0.93% to 8.31% with an average of 4.54% SOX2+ of all nuclei. These data confirm that SOX2+ cells can be present across PPGLs irrespective of tumour type, metastatic status and mutation status.

**Figure 1.**
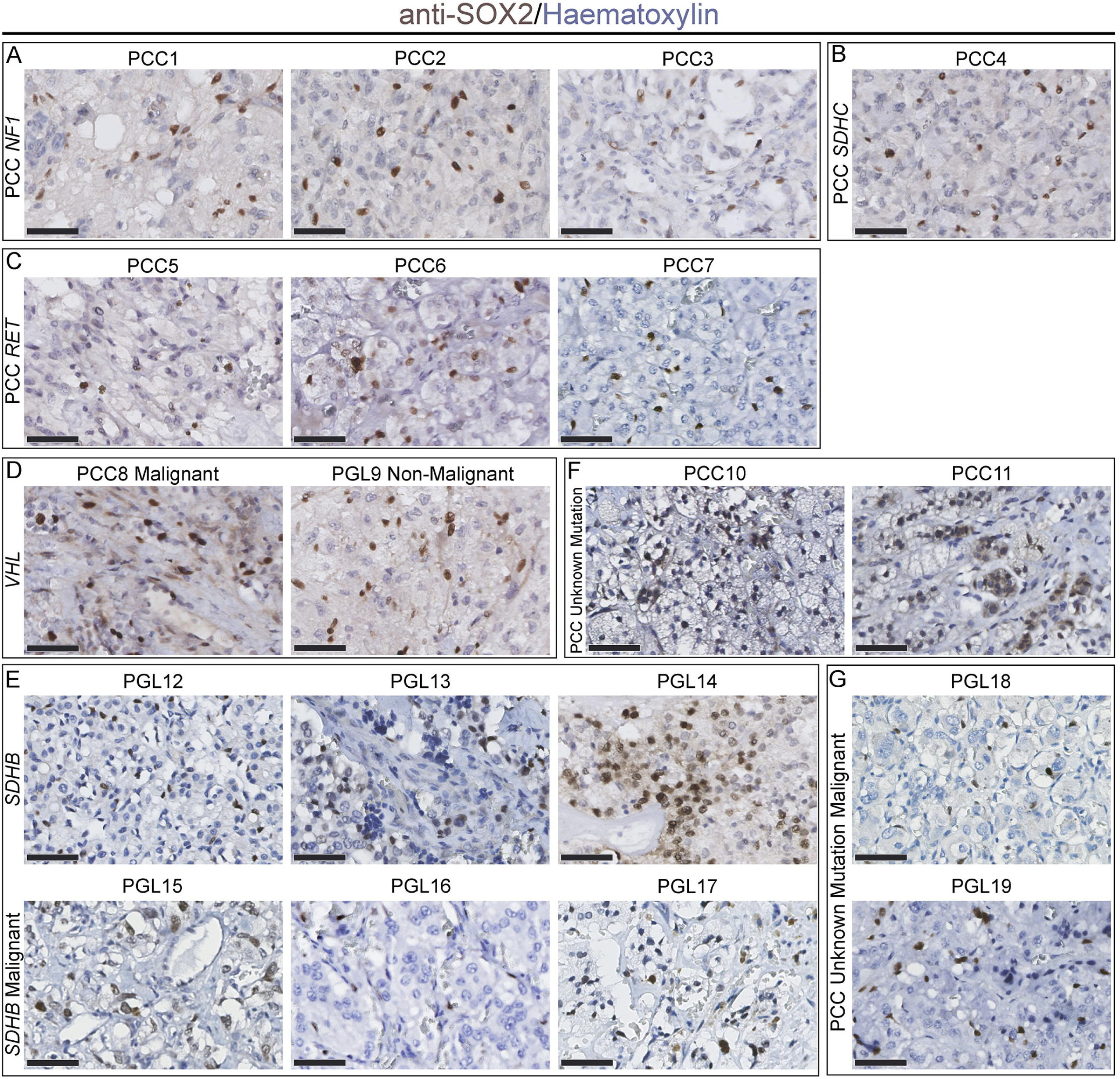
Immunohistochemistry using antibodies against SOX2 on FFPE sections of phaeochromocytomas and paragangliomas (PPGLs). In all panels, cells immunopositive for SOX2 are in brown, cells counterstained with haematoxylin. (A) Non-metastatic phaeochromocytomas PCC1 to PCC3 with mutations in NF1. (B) Non-metastatic phaeochromocytoma PCC4 with mutation in *SDHC*; (C) Non- metastatic phaeochromocytomas PCC5 to PCC7 with mutations in *RET* (D) Malignant phaeochromocytoma PCC8 and non-malignant paraganglioma PGL9 with mutations in *VHL* (E) Non- malignant paragangliomas PGL12 to PGL14 and malignant paragangliomas PGL15 to PGL17 with mutations in *SDHB* (F) Non-malignant phaeochromocytomas PCC10 and PCC11 with no known mutations. (G) Malignant phaeochromocytomas PCC12 and PCC13 with no known mutations. Scale bars 50µm.

**Table 1.**
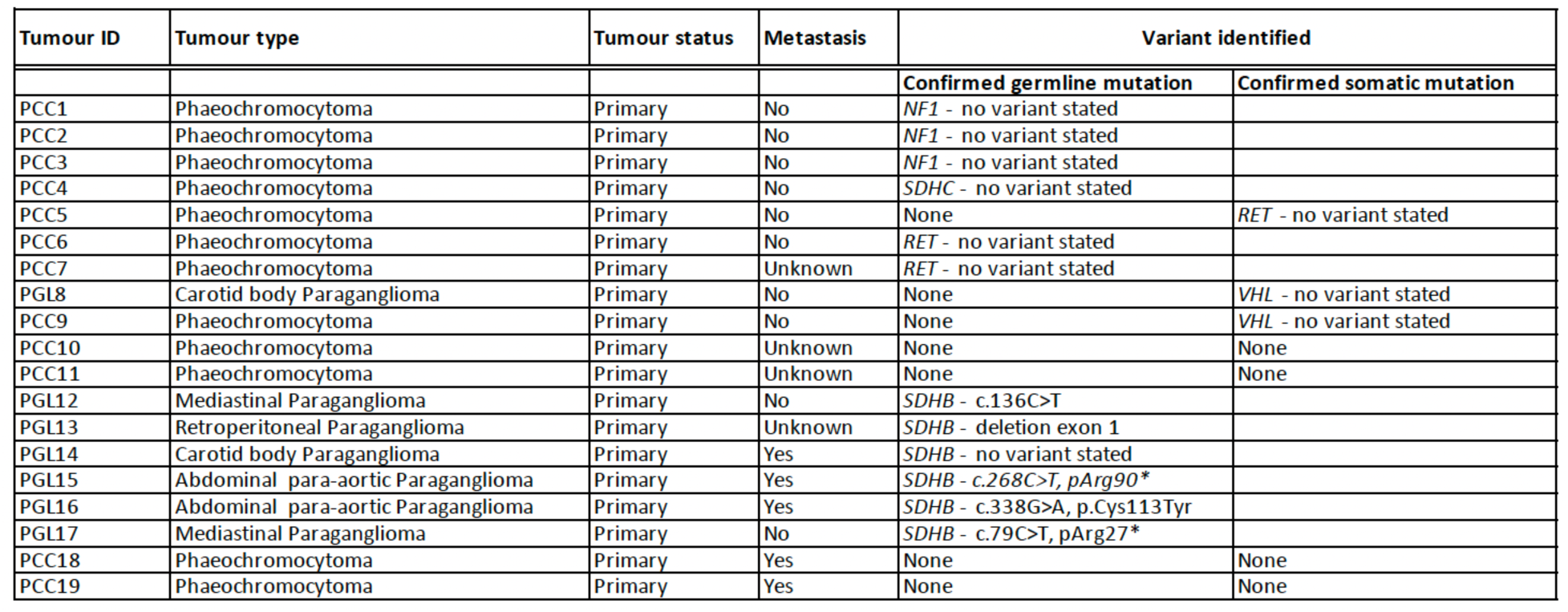
Details of tumour samples analysed by immunohistochemistry in Figure 1.

### Single-cell analysis of PPGLs identifies a second *SOX2*-expressing population of chromaffin cells in tumours

To examine the molecular profile of tumour cell populations, we took advantage of published single-nuclei datasets (10) including nine PGLs (PGL20, PGL21, PGL23 to PGL29) and one PCC (PCC22). Additionally, we carried out single-cell analysis on three fresh tumour samples (PCC30, PGL31, PGL32) and single-nuclei analysis on an additional four frozen samples (PGL33 to PGL36). Of the seven new samples, two were metastatic (PGL32, PGL33) and three had germline mutations in *SDHB* and two in *SDHD* (one metastatic). All tumour data used for transcriptomic analysis are shown in **Table 2**. For comparisons, we included two published single-nuclei datasets of normal adrenal tissue (Healthy1 and Healthy2) (15).

**Table 2.** Details of tumour samples used for transcriptomic analyses.

| Source | Tumour ID in original publication | Tumour ID in this publication | Tumour Type | Single Cell or Single | Tumour status | Metastasis | Metastasis site | Variant identified | SOX2+ cells |
| --- | --- | --- | --- | --- | --- | --- | --- | --- | --- |
| Flynn et al. 2025 | E019-P1 | PGL20 | Abdominal Paraganglioma | SN | Primary | No | NA | *SDHB* c.418G>T | Yes |
| Flynn et al. 2025 | E123-M1 | PGL21 | Abdominal Paraganglioma | SN | Metastasis | Yes | Bone | *SDHB* c.380T>G | No |
| Flynn et al. 2025 | E140-P1 | PCC22 | Phaeochromocytoma | SN | Primary | No | NA | *SDHB* c.380T>G | Yes |
| Flynn et al. 2025 | E143-M1 | PGL23 | Abdominal Paraganglioma | SN | Metastasis | Yes | Abdominal | *SDHB* c.268C>T | Yes |
| Flynn et al. 2025 | E146-P1 | PGL24 | Abdominal Paraganglioma | SN | Primary | No | NA | *SDHB* c.136C>T | Yes |
| Flynn et al. 2025 | E156-P1 | PGL25 | Bladder Paraganglioma | SN | Primary | No | NA | *SDHB* c.526G>T | Yes |
| Flynn et al. 2025 | E166-M1 | PGL26 | Abdominal Paraganglioma | SN | Metastasis | Yes | Abdominal | *SDHB* c.72+1G>T | Yes |
| Flynn et al. 2025 | E171-M1 | PGL27 | Abdominal Paraganglioma | SN | Metastasis | Yes | Chest wall - left rib 5-9 | *SDHB* c.736A>T | No |
| Flynn et al. 2025 | E197-M1 | PGL28 | Abdominal Paraganglioma | SN | Metastasis | Yes | Thoracic mediastinum lymph node | *SDHB* c.1 72del | Yes |
| Flynn et al. 2025 | E225-P1 | PGL29 | Abdominal Paraganglioma | SN | Primary | No | NA | *SDHB* c.190delG | Yes |
| This publication | NA | PCC30 | Phaeochromocytoma | SC | Primary | No | NA | None | Yes |
| This publication | NA | PGL31 | Abdominal Paraganglioma | SC | Primary | No | NA | *SDHB* c.268C>T p(Arg90Ter) | Yes |
| This publication | NA | PGL32 | Abdominal Paraganglioma | SC | Primary | Yes | Skeletal | None | Yes |
| This publication | NA | PGL33 | Carotid body Paraganglioma | SN | Primary | Yes | Pulmonary | *SDHD* c.337_340del p.(Asp113fs) | Yes |
| This publication | NA | PGL34 | Abdominal Paraganglioma | SN | Primary | No | NA | *SDHB* c.268C>T p.(Arg90*) | Yes |
| This publication | NA | PGL35 | Pelvic Paraganglioma | SN | Primary | No | NA | *SDHB* Exon 1 Deletion | Yes |
| This publication | NA | PGL36 | Carotid Body Paraganglioma | SN | Primary | No | NA | *SDHD* c.242C>T, p.(P81L) | Yes |
| Zethoven et al. 2022 | E240 | Healthy 1 | NA | SN | NA | No | NA | NA | Yes |
| Zethoven et al. 2022 | E243 | Healthy 2 | NA | SN | NA | No | NA | NA | Yes |

Our analysis identified nine clusters of cells, including B-cells, Chromaffin cells, Cortex, Endothelial cells, Fibroblasts, Macrophages, Mast cells, Sustentacular cells and T-cells (**Figure 2A**). Sample distribution split between the two healthy samples and 17 tumours revealed the Cortex cell cluster derives almost entirely from the healthy adrenal controls, with representation from both groups in all other clusters. Key markers of cell populations known from mouse data are plotted in **Figure 2B**; for chromaffin cells *HAND2*, *SLC18A1*, *TH* and *PHOX2A* and for the sustentacular population *SOX10*, *CDH19*, *S100B* and *SOX2*. The markers defining each cell population are listed in **Supplementary Table 1**. Top sustentacular markers included previously undescribed *PTPRZ1*, *KIRREL3*, *NKAIN3* and *CHST9* (**Figure 2C**). Proportions of all clusters in each of the datasets reveal variation in the sustentacular cells, with an over-representation of this cluster in PGL33 (31%) (**Figure 2D**). Comparison of the Chromaffin cell cluster of tumours from patients with metastatic disease (7 tumours) to those from tumours that had not metastasised at the time of analysis (10 tumours), identified factors potentially associated with metastasis. The top candidate metastasis genes were *GLCE*, *NELL1*, both involved in the tumour matrisome (16), *C17orf97* (LIAT1), *GHR* encoding the growth hormone receptor, known to play a role in cancer metastasis and chemoresistance (17, 18), *MND1*, involved in cell cycle regulation and DNA repair and a prognostic marker for several cancers (19–21), and *CREB5*, proposed to promote invasion and metastasis in colorectal cancer and glioma (22, 23). *EZH2*, a histone methyltransferase that promotes cancer progression (24, 25), as well as *PIMREG*, promoting aggressiveness through NF-κΒ and prognostic for several tumour types (26–28) are also in this group (**Figure 2E**). *EZH2*, *MND1* and *PIMREG* have been independently identified as associated with metastasis in *SDHx* tumours using bulk RNA-seq (15).

**Figure 2.**
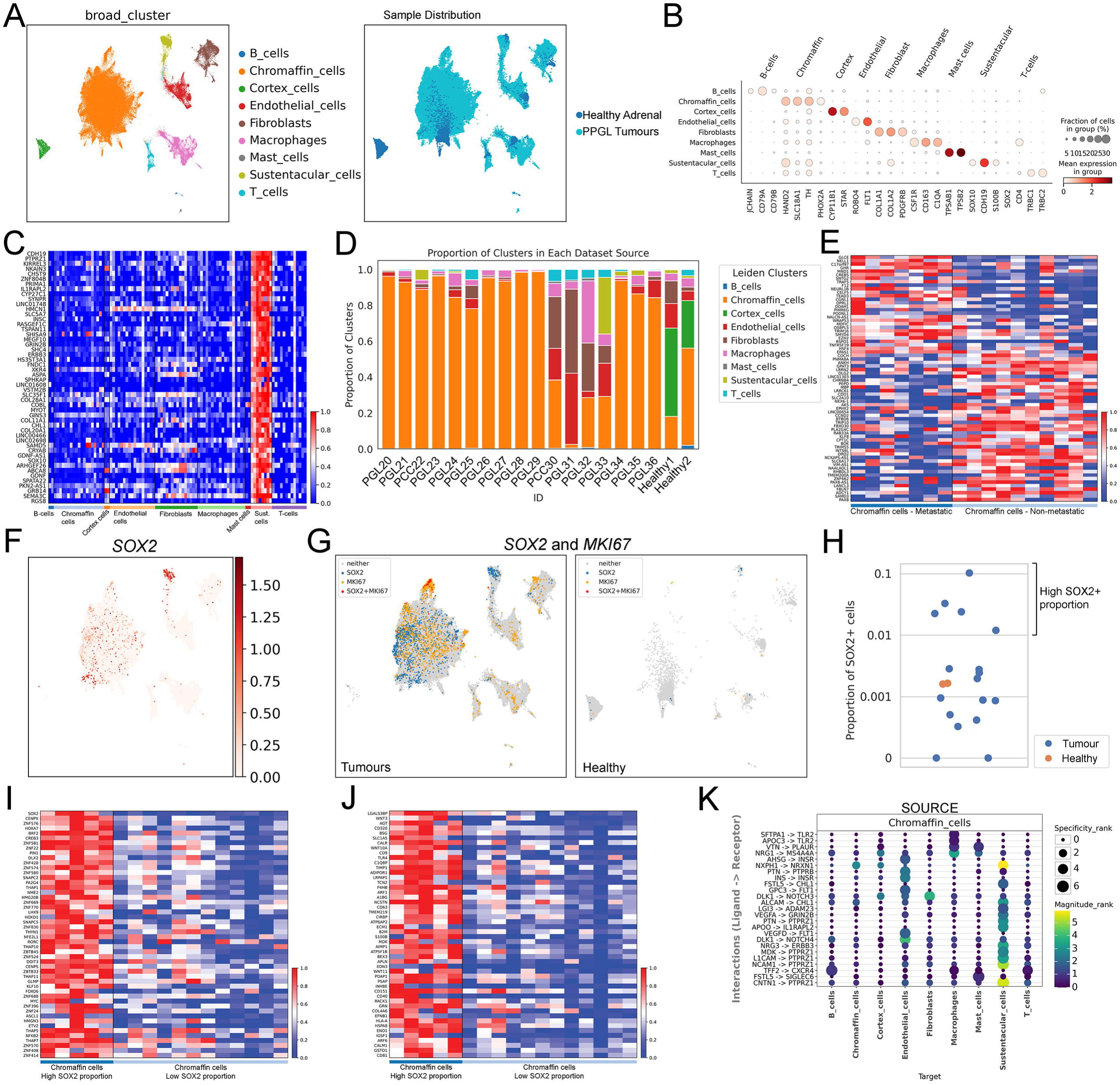
Single-cell analysis of PPGLs identifies expression of SOX2 in tumour chromaffin cells. (A) UMAP plot of the integrated dataset with each cell type (left) and patient group (right - healthy vs tumour) coloured separately. (B) Dotplot of key markers associated with the identified cell populations. (C) Heatmap of the top marker genes in sustentacular cells ordered by statistical significance from top to bottom. Colours correspond to gene-wise min-max normalised values. (D) Stacked barplot of cell type proportions in each sequenced sample; (E) Heatmap of top marker genes in chromaffin tumour cells comparing metastatic (left) to non-metastatic (right) samples. Genes were ranked by statistical significance from top to bottom for upregulated genes (for metastatic) and bottom to top for downregulated genes. Colours correspond to gene-wise min-max normalised values. (F) UMAP plot of normalised *SOX2* expression across all tumour cells, with the dots ordered by their expression values. (G) UMAP showing *SOX2* (blue), *MKI67* (cycling cells, yellow) and double-positive cells (red). Cells were classified as positive if they had at least a single transcript. (H) Strip plot of the proportion of *SOX2*+ cells in each sample. Distances on the Y-axis are on a log10 scale, and for visualisation purposes, datasets with 0% SOX2+ were plotted at Y=0.0001. Jitter has been added to the X-axis to reduce overlapping observations. (I) Heatmap of top transcription factor-encoding genes distinguishing chromaffin tumour cells in tumours with high SOX2+ cell proportion, from chromaffin tumour cells in tumours with low SOX2+ cell proportion, as determined with a 1% cut-off (see H). Genes are ordered by statistical significance from top to bottom. Colours correspond to gene-wise min-max normalised values. (J) Heatmap of genes associated with signalling, distinguishing chromaffin tumour cells in tumours with high SOX2+ cell proportion, from chromaffin tumour cells in tumours with low SOX2+ cell proportion, as determined with a 1% cut-off (see H). Genes are ordered by statistical significance from top to bottom. Colours correspond to gene-wise min-max normalised values. (K) LIANA+ dotplot with of the top 20 interactions where chromaffin cells are the source and ordered by specificity rank. Colours correspond to magnitude while dot size increases with specificity rank, as calculated by LIANA+.

Gene expression analysis of only the tumour samples confirms highest *SOX2* expression in the sustentacular cluster (**Figure 2F**). Unexpectedly, there was also widespread *SOX2* expression across the Chromaffin cell cluster. Projection of *SOX2* expression (blue) combined with *MKI67* (yellow) marking cycling cells, demonstrated that double-positive cells (red) are located in the Chromaffin cell cluster (**Figure 2G**, left). This confirms a population of cycling tumour cells that express *SOX2* but have chromaffin characteristics, unlike any population in healthy controls (**Figure 2G**, right). Healthy controls and 15 out of 17 tumours contained *SOX2*+ cells (**Supplementary** Figure 1). The proportion of *SOX2*-expressing cells varied across tumours, with a group of five tumours having a substantially higher proportion of *SOX2*+ cells than healthy controls (**Figures 2H**). Given the highly variable number of *SOX2*-expressing cells across samples, we decided to compare Chromaffin cells with higher proportion of *SOX2*+ cells to those with lower proportions or absence of *SOX2* (cut-off of 1%). Reassuringly, this analysis found the most differentially expressed transcription factor-encoding gene to be *SOX2*. Other highly ranked genes include *HOXA7*, whose protein product can function as an oncogene and contribute to malignancy (29–31), the proto-oncogene *BRF2* (32, 33) (**Figures 2I** and **Supplementary Table 2** for full list of significant factors). Our previous findings in mice demonstrated that SOX2+ cells have a major role in regulating cell proliferation of neighbouring cells through secretion of WNT6 (14). Comparative analysis of Chromaffin cells from tumours with a high SOX2 proportion versus those with lower/absent *SOX2*+ cell proportions, revealed differential expression of genes related to cell signalling. These include *LGALS3BP*, encoding a secreted factor involved in cancer progression (34), as well as the WNT protein-encoding genes *WNT3*, *WNT10A*, *WNT11* and *WNT6* (35–37) (**Figures 2J** and **Supplementary Table 2**). To further investigate the relationships of chromaffin cells with the other tumourigenic populations, we carried out ligand-receptor analysis framework (LIANA+) analysis (38). This determines the intercellular communication inference between chromaffin cells and the other populations. Interestingly, the strongest, in magnitude, predicted signalling from chromaffin cells is to sustentacular cells, primarily through adhesion molecules ASHG, CNTN1, NCAM1, L1CAM, ALCAM, as well as through NRG3, PTN and APOO. Signalling to endothelial cells and fibroblasts is predicted through DLK1 and to macrophages by NRG1 (**Figure 2K**). Taken together, these *in silico* data identify a malignant signature for PPGLs and support that tumour chromaffin cells aberrantly expressing SOX2, have distinct oncogenic gene expression and a secretory signature consistent with cancer promotion.

### PPGLs harbour cells that can be isolated in stem cell-promoting media and express SOX2 *in vitro*

We previously established a protocol for the isolation of stem cells from the mouse adrenal medulla (14). We sought to determine if these culture conditions can be used to isolate stem cells from human tissues. Foetal adrenal medullae at 12, 17 and 19 post-conception weeks (PCW) were immunostained using antibodies against SOX2. Positive cells were identified in all conditions (**Figure 3A**). Cells from an adrenal medulla at 19 PCW were dissociated into a single-cell suspension and plated in defined stem cell-promoting media (14). Cells adhered after 24 hours and colonies were visible at 96 hours (distinct by 72h), which could be expanded and passaged (**Figure 3B**). This stem cell isolation protocol was also applied to two fresh PPGL samples, a benign PCC with no known mutations (PCC30) and a metastatic PGL with a mutation in *SDHD* (PGL33); transcriptomic analyses confirmed *SOX2*-expressing cells in both tumours (**Supplementary** Figure 1). Colonies were obtained from both tumours *in vitro* (**Figure 3B**). Immunofluorescence staining of PCC30 cultures, confirmed the expression of SOX2 in expanded colonies (**Figure 3C**). Immunofluorescence against TH did not detect chromaffin cells in the control foetal sample (not shown) but did in PCC30 (**Figure 3C**). Isolated cells were capable of at least three passages and recovered one freeze-thaw cycle. These findings confirm the presence of a PPGL cell population with *in vitro* clonogenic properties, consistent with stem cell potential.

**Figure 3.**
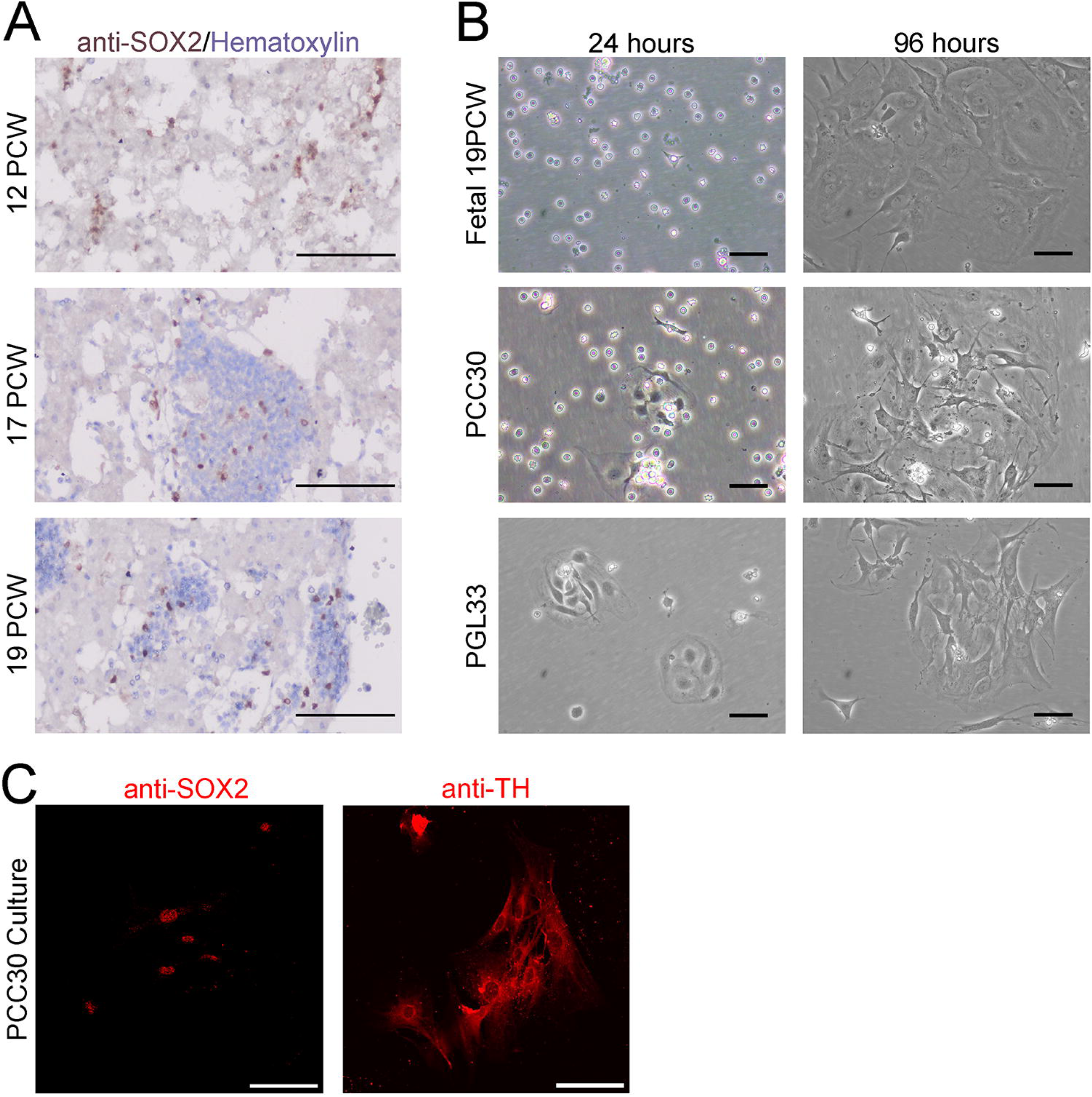
In vitro isolation and characterisation of PPGL-derived stem cells. (A) Immunohistochemistry confirming the presence of SOX2+ cells in foetal adrenal medullae at 12 post-conception weeks (PCW), 17PCW and 19PCW. SOX2 positive cells in brown, tissues counterstained with haematoxylin. Scale bars 100µm. (B) *In vitro* isolation of foetal adrenomedullary cells at 19PCW leads to the generation of adherent colonies from 3 days under defined stem cell-promoting media. Scale bars 50µm. *In vitro* isolation of adult adrenomedullary cells from PCC30 benign phaeochromocytoma with no known mutations and PGL33, malignant paraganglioma with a germline mutation in *SDHD*, under defined stem cell-promoting media. Adherent cells and colonies seen at 24 and 96 hours. Scale bars 50µm. (C) Immunofluorescence using antibodies against SOX2 and TH confirms protein expression in PCC30 adherent colonies after two weeks in culture. Scale bars 50µm.

### SOX2+ PPGL cells have tumour-initiating capacity *in ovo*

We next sought to determine if *in vitro*-isolated SOX2+ PPGL cells can regenerate and metastasise in a xenograft assay. Isolated and expanded human SOX2+ PPGL cells PCC30, PGL33, as well as control foetal adrenal cultures were dissociated into a single-cell suspension and labelled with CellTracker CMFDA label, which converts into a green, fluorescent dye intracellularly, allowing cell tracing without dye transfer (**Figure 4A**). Purified and labelled *in vitr*o-isolated stem cells, were seeded onto fertilised chick chorioallantoic membranes (CAMs) as a single-cell suspension (100,000 cells per CAM) and incubated for four days (see **Figure 4B** for schematic). CAM assays are established *in vivo* models for the study of tumour propagation, angiogenesis and metastasis (39–44). For each culture, grafting was carried out in a total of 10 eggs, over two independent experiments. Two eggs for PCC30, four eggs for PGL33 and four eggs for foetal controls survived and were analysed further. For PGL33, this led to the formation of tissue masses, presenting with green CMFDA label fluorescence (n=4) (**Figure 4C**). This was not observed in CAMs seeded with PCC30 (n=2 surviving eggs), or control foetal adrenal-derived stem cells (n=4 surviving eggs). Immunofluorescence using an antibody against human nuclear antigen (HNA) and immunohistochemistry using antibodies against SOX2 confirmed cells within the *de novo* formed masses are derived from seeded human PGL33 stem cells and continue to express SOX2 protein (**Figure 4D**). The lungs of each chicken embryo were harvested, fixed and processed for cryosectioning to enable analysis of possible metastasis through visualisation of green fluorescence. Detection of green CMFDA fluorescence in host chickens’ lungs revealed the presence of xenotransplanted, *SDHD*-mutant, PGL33-derived stem cells or their derivatives, confirming their potential to invade and metastasise (**Figure 4E**). Immunofluorescence staining using antibodies against SOX2 and TH on chicken lungs from PGL33 xenotransplantations demonstrated the presence of SOX2+ cells, TH+ chromaffin cells, and infrequent double-positive cells (**Figure 4F**). In summary, *in vitro*-isolated PPGL stem cells have tumour-initiating and metastatic capacity in a xenograft assay.

**Figure 4.**
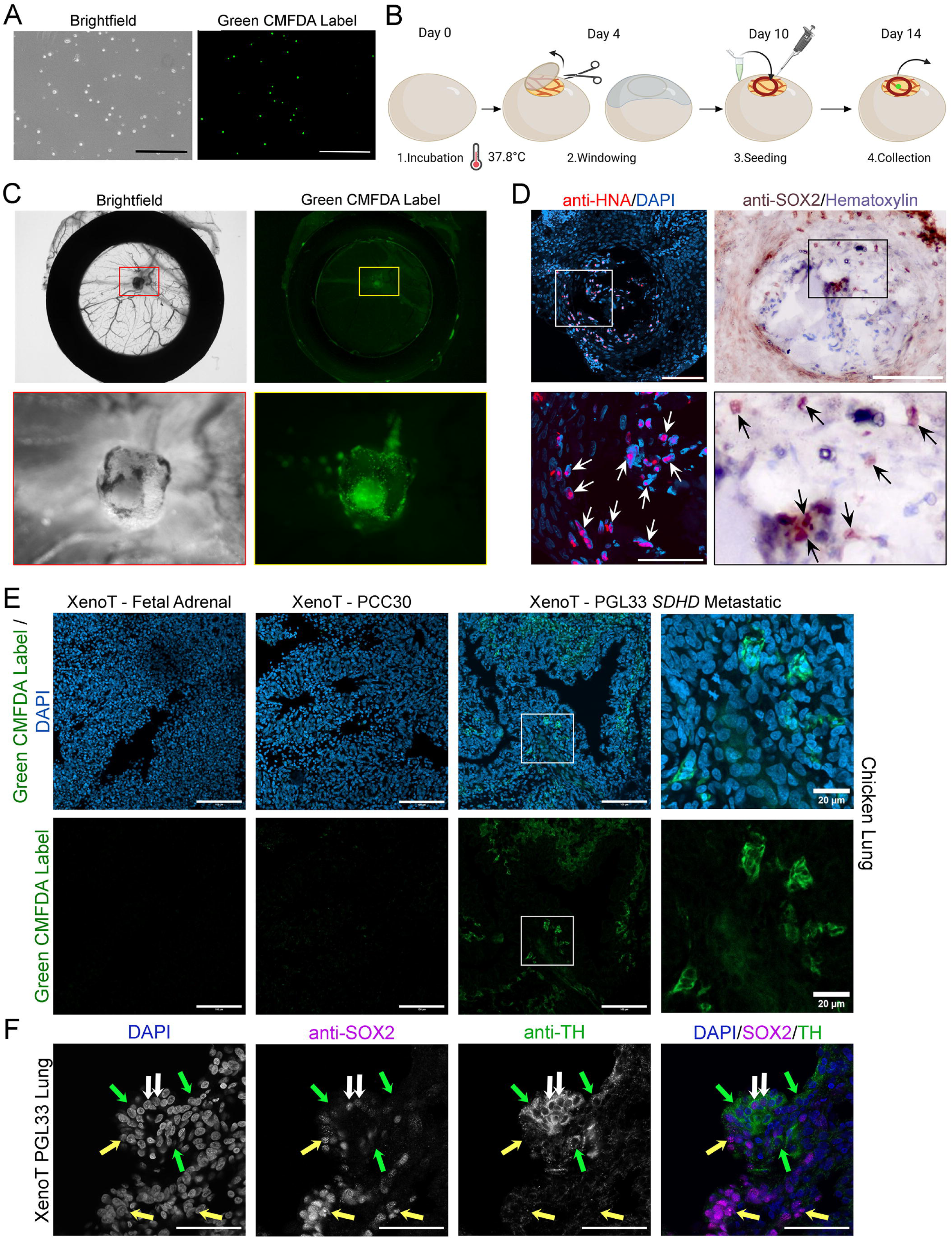
SOX2+ PPGL stem cells have tumour-inducing and metastatic capacity in ovo. (A) Single-cell suspension and labelling of foetal 19PCW cells with CellTracker green CMFDA label. Scale bars 500µm. (B) Pipeline for the *in vitro* isolation and *in ovo* transplantation of human SOX2+ cells (schematic created in BioRender.com). (C) Chick chorioallantoic membrane (CAM) xenograft of PGL33 after 4 days of incubation. Brightfield and UV fluorescence images of the dissected CAM, with a visible cell mass established on the vascularised CAM. The silicon O-ring is visible. Red and yellow boxes indicate the magnified regions shown below, depicting the mass. (D) Immunostaining on formalin-fixed, paraffin sections of the xenograft in C. Immunofluorescence using antibodies against human nuclear antigen (HNA, red)) confirming the presence of human cells in the mass. Nuclei counterstained with DAPI. Immunohistochemistry with antibodies against SOX2 (brown), confirming the presence of SOX2-expressing cells. Nuclei counterstained with haematoxylin. Scale bars 100µm. (E) Cryosections through the lungs of chicks where CAMs were successfully grafted with either foetal adrenal stem cell cultures, isolated stem cell cultures from PCC30 or isolated stem cell cultures from PGL33. Green fluorescent cells are detected in PGL33 cultures confirming metastasis. Nuclei counterstained with DAPI. Scale bars 100µm, except on right-hand panels where scale bars are 20µm. (F) Immunofluorescence staining on cryosections through chick lungs of a PGL33 xenograft with metastasis, using antibodies against SOX2 (magenta) and TH (green). Nuclei counterstained with DAPI (blue). The greyscale images of single channels are shown for signal clarity. Yellow arrows indicate SOX2 positive cells, green arrows indicate TH positive cells and white arrows indicate SOX2;TH double-positive cells. Nuclei counterstained with DAPI. Scale bars 50µm. XenoT = xenotransplanted.

## DISCUSSION

Identifying the cell of origin in tumours and cancers is critical for developing accurate experimental models of tumourigenesis, discovering prognostic markers, and the design of targeted therapies. The paradigm of cancer stem cells or tumour-initiating cells contributing directly to tumour formation and maintenance has been established in various solid tumours (45–51) We recently identified a neural crest-derived postnatal stem cell population expressing *SOX2*. Here, we demonstrate that *SOX2+* cells are consistently present in PPGLs, regardless of tumour location, malignancy status, or underlying genetic alterations. *In silico* analysis of 17 PPGL single-cell RNA sequencing datasets corroborates widespread *SOX2* expression in all but two tumours and defines a robust molecular signature linked to metastatic potential. In addition to *EZH2*, *MND1* and *PIMREG*, which are known promoters of cancer progression and have previously emerged as candidate metastatic markers in PPGL (15), we report additional novel candidates associated with metastatic behaviour. One particularly promising candidate is cyclic adenosine monophosphate (cAMP)-response element-binding protein 5 (*CREB5*). In colorectal cancer, *CREB5* can promote invasion and metastasis through activation of the receptor tyrosine kinase *MET*, and is associated with an unfavourable prognosis in multiple cancers including hepatocellular carcinoma, glioma, breast, prostate, and epithelial ovarian (22, 23). These candidates require functional validation to determine the true drivers of metastasis, which will subsequently guide the identification of actionable therapeutic targets. Future work encompassing larger tumour cohorts will better resolve these drivers and aid in prognostic predictions.

Stem cells have been implicated in cancer relapse and metastasis (52, 53) and can regenerate tumours when grafted (54). This study provides compelling evidence that PPGL-derived *SOX2*+ cells possess tumour-initiating capacity, with the ability to self-renew *in vitro*, regenerate and metastasise in xenotransplantation assays - key hallmarks of cancer stem cells (46). We identified two distinct *SOX2*+ populations within PPGLs; a sustentacular population, also present in healthy controls, and a tumour-specific subpopulation of chromaffin cells, echoing earlier histological observations (13). The presence of double-positive cells (expressing both *SOX2* and chromaffin markers) exclusively in tumours is particularly intriguing; these cells may be a unique stem cell of the tumours, since these are absent in normal adrenal tissue. Such cells may arise either from re-activation of a stem-like transcriptional programme in chromaffin cells, or from aberrant differentiation of sustentacular cells whilst they retain *SOX2* expression instead of downregulating this gene as in normal differentiation. Currently, it remains unclear whether metastasis in xenografts originates from *SOX2*+ tumour chromaffin cells or the *SOX2*+ sustentacular stem cell population.

Supporting a direct role in tumour growth, *SOX2*+ tumour chromaffin cells actively proliferate, unlike the more quiescent *SOX2*+ sustentacular cells. Moreover, they display a distinct transcriptomic signature that is enriched in genes promoting tumourigenesis and cancer progression. Our prior work demonstrated that WNT6 secretion by adrenomedullary stem cells regulates chromaffin cell proliferation (14). In the present study, the expression of several *WNT* genes, including *WNT6* and additional pro-tumourigenic factors by *SOX2*+ tumour chromaffin cells suggests a possible paracrine contribution to tumourigenesis, mirroring mechanisms observed in other cancers (55–61).

Overall, our findings establish that PPGLs contain cells with stem-like properties that can be isolated, expanded, and transplanted, and have the capacity to propagate tumours. These results provide the first functional evidence suggesting that *SOX2*+ cells in PPGLs possess the potential to serve as tumour cells of origin.

## Acknowledgements

We thank the King’s College London Biological Services facilities, the Genomics Research Platform, R&D Department, Guy’s and St. Thomas’ NHS Trust. This research was supported by samples of the BioBank Dresden as a core facility of the Medical Faculty Dresden/Technical University Dresden and National Center of Tumor Diseases Dresden resource.

