## Supplementary figures and images for "Phaeochromocytomas and paragangliomas harbour tumour-initiating SOX2+ stem cells"

### Supplementary Figure 1

#### Supplementary Figure 1

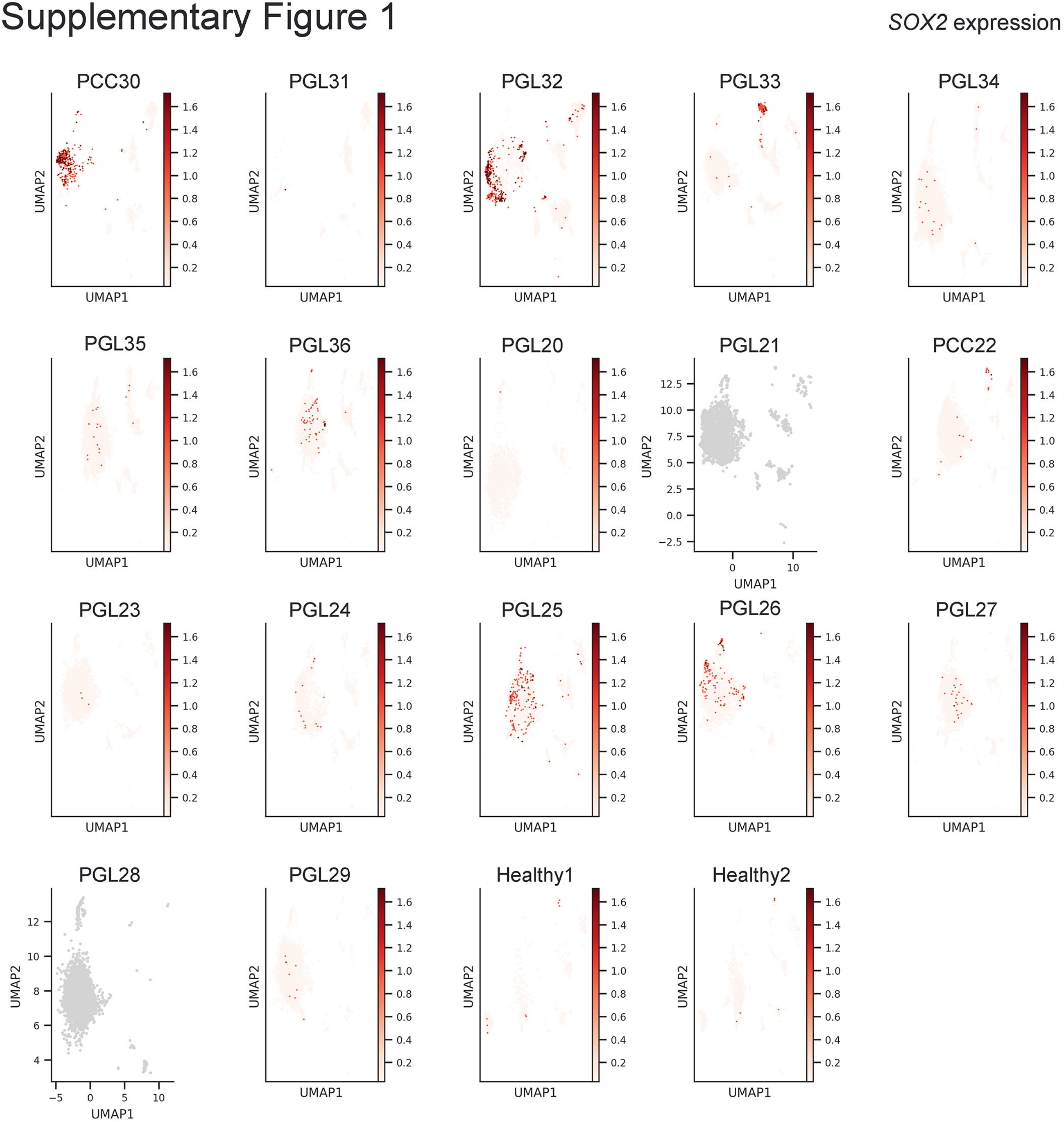
